# Phylogenomics of 8,839 *Clostridioides difficile* genomes reveals recombination-driven evolution and diversification of toxin A and B

**DOI:** 10.1101/2020.07.09.194449

**Authors:** Michael J. Mansfield, Benjamin J-M Tremblay, Ji Zeng, Xin Wei, Harold Hodgins, Jay Worley, Lynn Bry, Min Dong, Andrew C. Doxey

## Abstract

*Clostridioides difficile* is the major worldwide cause of antibiotic-associated gastrointestinal infection. A pathogenicity locus (PaLoc) encoding one or two homologous toxins, toxin A (TcdA) and toxin B (TcdB) is essential for *C. difficile* pathogenicity. However, toxin sequence variation poses major challenges for the development of diagnostic assays, therapeutics, and vaccines. Here, we present a comprehensive phylogenomic analysis 8,839 *C. difficile* strains and their toxins including 6,492 genomes that we assembled from the NCBI short read archive. A total of 5,175 *tcdA* and 8,022 *tcdB* genes clustered into 7 (A1-A7) and 12 (B1-B12) distinct subtypes, which form the basis of a new method for toxin-based subtyping of *C. difficile*. We developed a haplotype coloring algorithm to visualize amino acid variation across all toxin sequences, which revealed that TcdB has diversified through extensive homologous recombination throughout its entire sequence, and formed new subtypes through distinct recombination events. In contrast, TcdA varies mainly in the number of repeats in its C-terminal repetitive region, suggesting that recombination-mediated diversification of TcdB provides a selective advantage in *C. difficile* evolution. The application of toxin subtyping is then validated by classifying 351 *C. difficile* clinical isolates from Brigham and Women’s Hospital in Boston, demonstrating its clinical utility. Subtyping partitions TcdB into binary functional and antigenic groups generated by intragenic recombinations, including two distinct cell-rounding phenotypes, whether recognizing frizzled proteins as receptors, and whether it can be efficiently neutralized by monoclonal antibody bezlotoxumab, the only FDA-approved therapeutic antibody. Our analysis also identifies eight universally conserved surface patches across the TcdB structure, representing ideal targets for developing broad-spectrum therapeutics. Finally, we established an open online database (DiffBase) as a central hub for collection and classification of *C. difficile* toxins, which will help clinicians decide on therapeutic strategies targeting specific toxin variants, and allow researchers to monitor the ongoing evolution and diversification of *C. difficile*.

## Introduction

*Clostridioides difficile* (formerly *Clostridium difficile*) is a diverse group of Gram-positive sporeforming anaerobic bacteria [1]. Toxigenic strains have become important opportunistic pathogens to humans. Their spores are widespread and can colonize human and animal colons after disruption of the gut microflora, most notably due to antibiotic treatment. *C. difficile* infection (CDI) results in a range of symptoms from self-limiting diarrhea to severe pseudomembranous enterocolitis and death [2–7]. It is the most frequent cause of healthcare-associated gastrointestinal infections across developed countries worldwide [2–5,8].

Ribotyping (RT), which compares intergenic spacers between ribosomal RNA genes, is widely utilized to categorize *C. difficile* linages [5,9]. Various other methods including multilocus sequence typing based on allelic variation of housekeeping genes and whole genome sequencing analysis have also been adopted to further discriminate strains [5,9–13]. Phylogenetic analyses revealed a growing diverse population [1,14–16]. In recent years, there is an emergence and spreading of various epidemic hypervirulent strains such as the RT027 clonal lineage, which first caused outbreaks in 2000-2003 in North America and is associated with increased disease severity and mortality [17–20]. RT078 is an emerging hypervirulent lineage which is also the dominant type found in domesticated animals [21,22]. There are also geographic differences, for instance, RT017 has become a dominant lineage in Japan and Korea [23].

The major virulence factors in toxigenic *C. difficile* strains are two homologous large protein toxins, TcdA (~300 kDa) and TcdB (~270 kDa) [24–27]. Nontoxigenic *C. difficile* strains without these toxins exist and can colonize humans and animals, but do not cause diseases [28]. TcdA and TcdB share overall ~66% sequence similarity and belong to the large clostridial toxin (LCT) family, which include TcsH and TcsL in *Paeniclostridium sordellii*, Tcnα in *Clostridium novyi*, and TpeL in *Clostridium perfringens* [5,6,8,9,24,25,29–31]. TcsH and TcsL can be considered orthologs of TcdA and TcdB, respectively, with TcsH sharing ~77% sequence identity with TcdA and TcsL sharing ~76% identity with TcdB [32] (Fig. S1). TcdA and TcdB share a protein domain architecture consisting of an N-terminal glucosyltransferase domain (GTD), followed by a cysteine protease domain (CPD), an intermingled membrane translocation delivery domain and receptor-binding domain (DRBD), and a large C-terminal combined repetitive oligopeptides domain (CROPs) (Fig. S1). After binding, endocytosis, and translocation across endosomal membranes into the cytosol of host cells, these toxins glucosylate and inactivate host Ras/Rho family of small GTPases, leading to disruption of the actin cytoskeleton, cell rounding, and ultimately cell death [33].

TcdA and TcdB were first identified, characterized and sequenced in the 1980s [34–38] and the toxin sequences from a reference strain (VPI10463) have been widely used as the standard in diagnostic and therapeutic development. However, sequence variations in the toxin genes exist across *C. difficile* strains and could affect receptor-binding specificity, preferences toward distinct small GTPases, overall toxicity, and antigenicity. For instance, strains such as R20291 (belonging to RT027) produces a TcdB variant with ~8% of residue differences from the reference TcdB, which exhibited a significant impact on its immunogenicity: mice immunized with the reference TcdB developed resistance to the same TcdB, but all died when challenged with this variant TcdB [39], and several antibodies raised against the reference TcdB, including the FDA approved therapeutic antibody bezlotoxumab, either do not recognize or have lower efficacy against this TcdB variant [39–41]. Furthermore, this TcdB variant also lacks the ability to recognize frizzled (FZD) proteins, which are one of the major receptors for the reference TcdB, due to residue changes at the FZD-binding interface [40,42–45].

These toxin variations pose a significant challenge for developing effective broad-spectrum diagnostic assays, therapeutic antibodies, and vaccines. Understanding variations in toxins is a key step to address this challenge and may also reveal their potential evolutionary paths and functional differences. A toxinotyping method has been previously developed utilizing PCR-based amplification of toxin gene fragments and analyzing polymorphism with restriction enzyme digestions, which can distinguish over 34 toxinotypes [46,47]. Although toxinotyping highlights the variation among toxin genes, it lacks the resolution to understand the molecular basis for diversification of toxins and sequence-function relationships.

Rapid growth of genomic sequencing of *C. difficile* strains in recent years provides an opportunity to analyze and categorize the diversification of TcdA and TcdB with single residue resolution. Here we performed a comprehensive analysis of nearly all available *C. difficile* TcdA and TcdB sequences, including assembly and analysis of 6,492 new genomes, with the goal to 1) build a comprehensive central database of *C. difficile* toxin sequences; 2) better understand the mechanisms underlying TcdA and TcdB diversification; and 3) develop a system to classify TcdA and TcdB into subtypes that allow clinicians and researchers to categorize and predict functional-immunological variations of any future sequenced *C. difficile* isolates.

## Results

### Collection of TcdA and TcdB sequences across 8,839 C. difficile genomes

To build a comprehensive database of TcdA and TcdB sequences, we combined data from NCBI GenBank and the NCBI short-read archive (SRA). From 2,347 *C. difficile* genomes in GenBank, we identified an initial set of 1,633 *tcdA* and 2,028 *tcdB* genes. We then developed a computational pipeline for automated retrieval of *C. difficile* genomes from the SRA, *de novo* genome assembly, genome annotation, and extraction of *tcdA* and *tcdB* genes (see Methods). Using this pipeline, we assembled the genomes of 6,492 *C. difficile* isolates and identified an additional 3,542 *tcdA* and 5,994 *tcdB* genes (Table 1). Combining both sources, we identified 5,175 TcdA and 8,022 TcdB encoding sequences.

**Table 1.** Assembled *C. difficile* genomes from the NCBI SRA and associated statistics.

| Property | Statistic (mean +/- SD) |
| --- | --- |
| Number of samples | 6,492 |
| Assembly length | 4.2759 +/- 0.019 Mb |
| Number of contigs | 403.6 +/- 544 |
| GC content | 28.33 +/- 4.14 % GC |
| Mean contig length | 62.68 +/- 46.97 Kb |
| Contig N50 | 905,900.13 +/- 865,085 |
| Contig N90 | 298,748 +/- 52,843.68 |

We then carried out alignments of all toxin protein sequences. The TcdB alignment covered the entire sequence (1-2366), with 712 (30%) of the positions showing variations across all domains. The TcdA alignment possessed much lower variation than TcdB within the 1-1874 region as it had only 168 (9%) variable sites, but its CROPs domain (1831-2710) contained an extremely high degree of variation in the number length of repeats: from 3 repeats in the shortest variant to 45 in the longest variant, and 32 in the reference TcdA variant from VPI 10463 (Fig. S2). This is likely generated by homologous recombination due to the repetitive nature of this region. The CROPs domain is composed of long repeats (LRs) of ~30 residues and short repeats (SRs) of ~19-24 residues [27]. The CROPs domain in TcdA is not only repetitive at a protein sequence level, but also showed a high degree of repetitiveness at a DNA level, whereas the repetitiveness of the CROPs domain in TcdB is largely limited to the protein level [48,49], which may account for frequent recombination in TcdA-CROPs but not in TcdB-CROPs. It is also important to note that the repetitive nature of the TcdA-CROPS region at the DNA level may result in assembly errors, which may inflate the apparent variation in this region.

### Classifying TcdA and TcdB into subtypes

In total, there were 116 unique TcdA protein sequences and 212 unique TcdB protein sequences. We then clustered these sequences into distinct subfamilies (“subtypes”) using average linkage hierarchical clustering (see Methods). Analysis of TcdB is based on full-length sequences, but TcdA is limited to the 1-1874 region to avoid the highly variable CROPs domain. In addition, we also included TcsH and TcsL sequences in our analysis. Clustering produced 7 distinct TcdA subtypes which we labeled A1-A7, and 12 distinct TcdB subtypes which we labeled B1-B12 (Fig. 1). B1-B4 were ordered consistent to match previous literature [50–52], and additional subtypes are ranked based on total frequency of occurrence in GenBank and NCBI-SRA. Each unique sequence was then further numbered following a period within its subtype (e.g. B1.1, 1.2, 1.3, etc.). Sequences within the same TcdA and TcdB subtype demonstrate strong pairwise similarities, and weak similarities between subtypes (Fig. 1a, 1d). Quantitative analysis revealed that thresholds of 99.4% (TcdA) and 97% (TcdB) can be used to effectively assign toxin sequences to these subtypes (Fig. S3). We then selected one representative sequence for each subtype and carried out phylogenetic analysis and pairwise comparison. TcdA subtypes A1 to A6 possess higher similarities (>97.9%) and clustered together, with A7 forming a divergent lineage (Fig. 1b, 1c). A7 is a unique sequence with only 85.3% to 85.6% identity to others (Fig. 1c). The entire TcdA family was further outgrouped by TcsH as expected (Fig. 1b). TcdB also formed a monophyletic family that was outgrouped by TcsL and a second lineage of TcsL-related proteins (Fig. 1e). TcdB subtypes can be subdivided into three groups, one including B3, B4, B7, B8, a second including B2, B9, B10, B11, and a third including B1, B5, B6, and B12 (Fig. 1e). The lowest identity among TcdB subtypes is 85.3% (between B7 and B12, Fig. 1f). A7, B10, B11, and B12 represent rare divergent subtypes recently reported: A7 is in strain RA09-70, which does not express TcdB [53]; B10, B11, and B12 were identified recently from strains CD10-165, CD160, and 173070, respectively [53], and all three strains lack TcdA.

**Figure 1.**
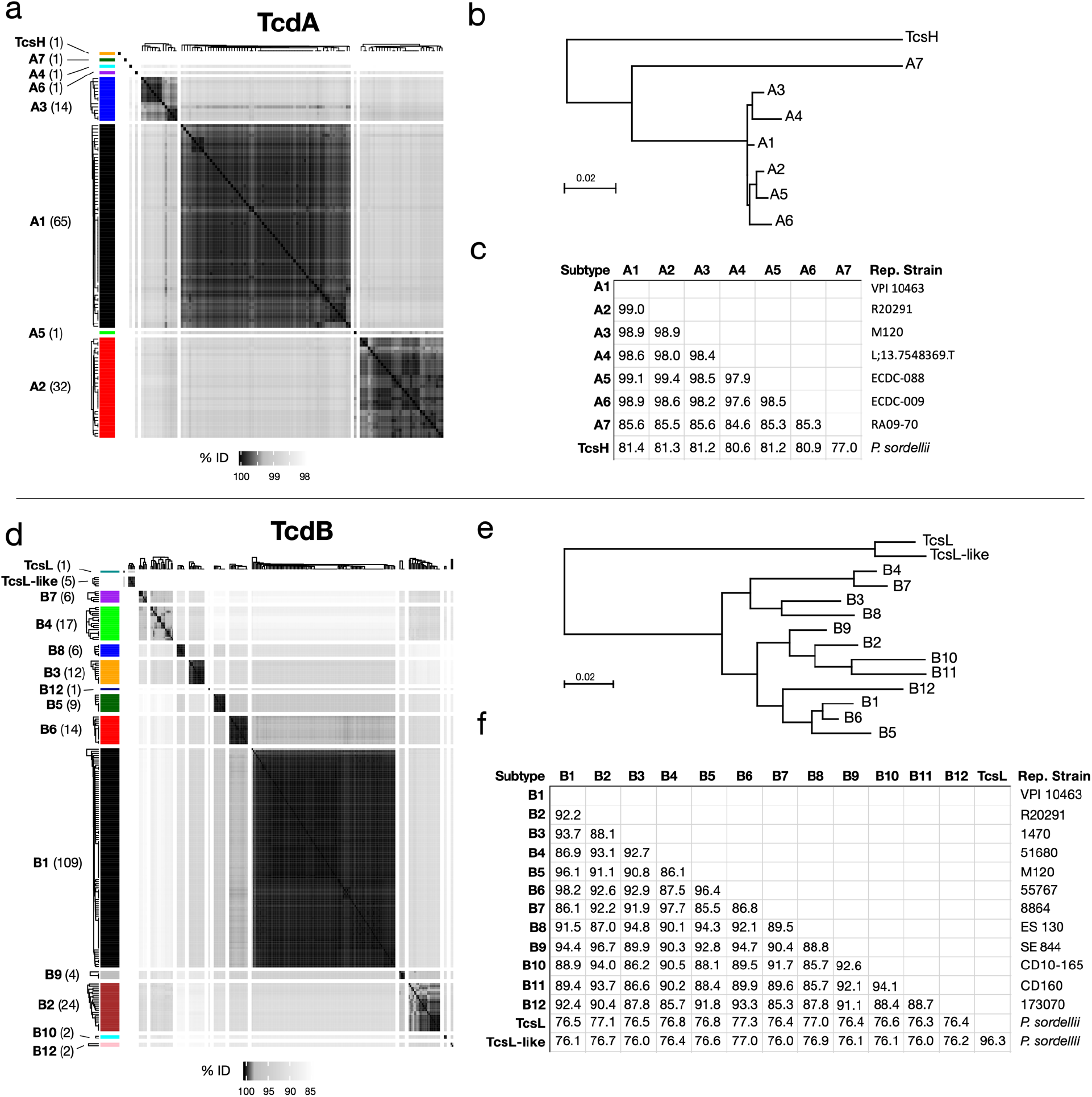
Clustering of TcdA and TcdB sequences derived from NCBI GenBank and SRA into subtypes. (**a**) Hierarchical clustering of TcdA sequences, split into 8 groups. (**b**) Neighbor-joining phylogenetic tree of representative sequences of each TcdA subtype. (**c**) Percentage identities between representative sequences. (**d**) Hierarchical clustering of TcdB sequences, split into 14 groups. (**e**) Neighbor-joining phylogenetic tree of representative sequences of each TcdB subtype. (**f**) Percentage identities between representative sequences. Hierarchical clustering was performed using the hclust() function in R, and cluster definitions were selected based on strong within-cluster sequence similarities and weak between-cluster similarities, as demonstrated visually and quantitatively. The reference strains (VPI 10463 and strain 630) are associated with TcdA group A1 and TcdB group B1a. The hypervirulent ribotype 027 strains such as R12087 and R20291 are associated with TcdA group A2 and TcdB group B2. Also included are the homologs of TcdA and TcdB (TcdH and TcdL, respectively) from *P. sordellii*, which expectedly exhibit the highest divergence from other groups. The datasets include TcdA and TcdB sequences from the NCBI GenBank as well as an additional 125 sequences assembled from the SRA.

By mining unassembled *C. difficile* genomes from the SRA, we were able to discover 125 TcdA and TcdB protein sequences that were not represented in GenBank. Most novel toxin variants clustered into subtypes A1 (N = 25), B1 (N = 52), and subtypes A2 (N = 10) and B2 (N = 12) (Table S1). However, three highly divergent TcdA variants identified from SRA datasets formed new subtypes not represented in GenBank. These include subtypes A4 from strain ECDC-088 (SRS1486236), A5 from strain ECDC-009 (SRS1486256), and A6 from strain L;13.7548369.T (SRS1486661), all of which are clinical isolates. All three of these strains contained truncated/partial TcdB variants which represent putative pseudogenes.

To link our subtyping with known clinical *C. difficile* strains, we manually curated subtype assignments for a set of 63 *C. difficile* strains selected from the literature, which covers known toxinotypes, and compared subtypes with toxinotypes, ribotypes, and whether the strain produces the third toxin known as *C. difficile* transferase toxin (CDT) (Table S2). The majority express an A1/B1 subtype combination and include reference strains 630 and VPI 10463 that express the widely used standard TcdA and TcdB sequence (defined as A1.1 and B1.1, Table S2). The group that expresses a combination of A2/B2 is the second largest and includes hypervirulent RT027 strains R12087 and R20291. The group expressing A3/B5 include strains (e.g. M120 and NAP07) classified as RT078. Subtype B3 is mainly expressed in strains (e.g. 1470) belonging to RT017, which lacks TcdA. Other pairings in the table include A2/B9, A3/B6, A2/B4, and A1/B3. The table includes many strains that do not express functional TcdA, which can express B1, B2, B3, B4, B5, B7, B8, B10, B11, or B12; one strain that only expresses TcdA but not TcdB (A7 in RA09-70); four strains that only express CDT; and one strain (SLO037) that expresses none of the three toxins. This table represents a small portion of *C. difficile* strains and a full list of a total 1,640 *C. difficile* strains from the NCBI database with their toxin subtypes noted is included as Table S3.

In general, phylogenetic subtyping of *C. difficile* toxins correlated well with previously identified toxinotypes, but at greater resolution by analyzing TcdA and TcdB separately (Table S2, see Discussion). There was less congruence with ribotypes, however, as different subtypes were found in the same ribotype strains, and the same subtype was found in different ribotype strains. Therefore, neither toxinotype nor ribotype were able to accurately categorize toxins based on phylogenetic relationships (Table S2). Subtyping was capable of capturing the full phylogenetic diversity of TcdA and TcdB available in previously known and new strains.

### Distribution of toxin subtypes across the C. difficile phylogeny

To evaluate the phylogenomic distribution of toxin subtypes across *C. difficile*, we constructed a whole-genome based phylogeny of 1,934 complete *C. difficile* genomes based on 14,194 SNP positions across 88 conserved marker genes (Fig. 2a, Table S3) (see Methods). The genome tree is highly consistent with known phylogenetic relationships, as the previously identified clades 1-5 are represented by distinct lineages [14] (Fig. 2a). Two of the three divergent environmental lineages C-I and C-II are also present as divergent branches (Fig. 2a).

**Figure 2.**
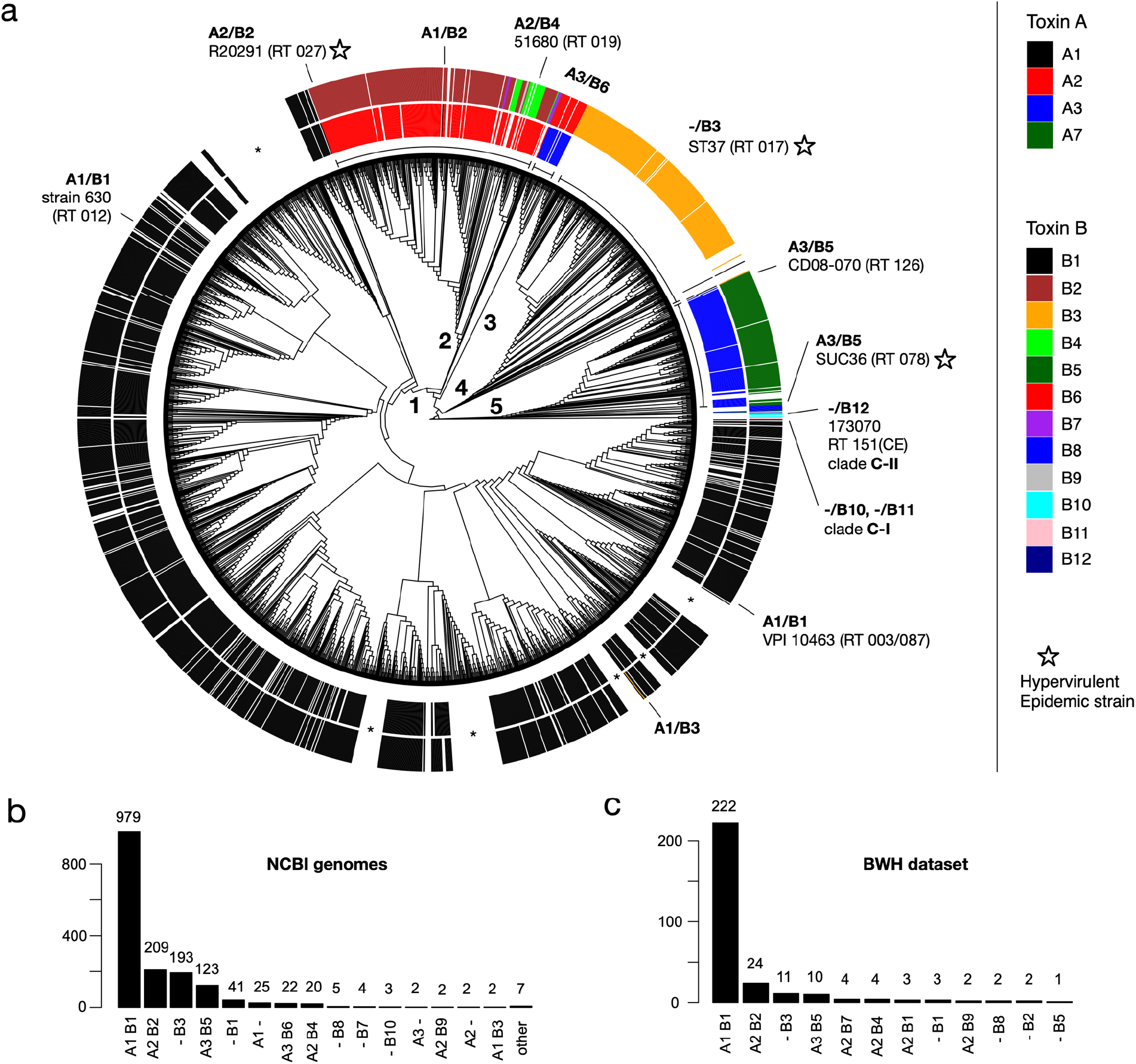
Toxin subtypes across the *C. difficile* phylogeny and occurrence of subtypes in a clinical CDI cohort. (**a**) TcdA (inner ring) and TcdB (outer ring) subtypes mapped onto a tree of 1934 *C. difficile* genomes. The genome tree is derived from the NCBI and is based on clustering of all-by-all genome BLAST scores. Lineages corresponding to previously identified *C. difficile* PaLoc clades (1 – 5) are labeled numerically. PaLoC clade 1 was subdivided into four sublineages labeled 1a-1d. Selected clinically relevant strains are shown on the tree, with hypervirulent/epidemic outbreak strains indicated by stars. Asterisks indicate lineages without toxin genes. (**b**) Frequency of toxin subtypes detected in 1,934 representative, complete *C. difficile* genomes from NCBI/GenBank. A total of 1,640 (84.8%) *C. difficile* strains contained TcdA and/or TcdB, while 294 (15.2%) were toxin deficient. (**c**) Frequency of toxin subtypes detected in a CDI clinical cohort from Brigham and Women’s Hospital (BWH). The total dataset contained 351 *C. difficile* genomes derived from infected patients. Of these, 289 (82.3%) contained toxin genes, and 62 (17.7%) were toxin deficient.

A total of 1,640 (84.8%) *C. difficile* strains were found to encode TcdA and/or TcdB, while the remainder (294, 15.2%) lack toxin genes. The predicted toxin subtypes across the *C. difficile* genome tree demonstrate strong clade associations, and therefore are highly congruent with strain phylogenetic relationships. The congruency between subtype and phylogeny provides further support for our toxin classification (Fig. 2a). For example, subtype A1/B1 which includes reference strains 630 and VPI 10463 is most common among toxin-containing strains (979, 59.7%) and associated with clade 1 (Fig. 2b). A2/B2 was second most common and associated with clade 2, A3/B6 with clade 3, -/B3 with clade 4, and A3/B5 with clade 5. Also prevalent were types -/B1, A1/-, and A2/B4 (Fig. 2b). Deviations from the A1/B1 toxin type are often associated with the emergence of numerous hypervirulent and epidemic outbreak strains such as A2/B2 (RT027), A3/B5 (RT078), and -/B3 (RT017) (Fig. 2a). Interestingly, the highly divergent environmental lineages encode the highly divergent TcdB subtypes B10 and B11 (C-I) and B12 (C-II) (Fig. 2a, Table S2). This is consistent with an early divergence of B10-B12 in *C. difficile* evolution, predating the emergence of TcdB subtypes found in the other clinical strains.

Interestingly, we also observed rare lateral transfer events involving only one of the two toxin genes to create hybrid strains containing new subtype combinations. Examples include the spontaneous emergence of an A1/<u>B3</u> strain within clade 1, and the emergence of an <u>A1</u>/B2 strain in clade 2 (Fig. 2a). Thus, through lateral transfer and homologous recombination, subtype B3 has likely replaced B1 in a clade 1 strain, and subtype A1 has likely replaced A2 in a clade 2 strain. Furthermore, we observed many independent clades containing *tcdA*-/*tcdB*-*C. difficile* strains (e.g., see six lineages marked by asterisks in Fig. 2a). This is consistent with previously reported “defective” toxin clades [54], and indicates numerous independent losses of the pathogenicity locus throughout *C. difficile* evolution.

### Toxin subtyping of an independent dataset of clinical C. difficile isolates

As an independent test dataset for our toxin subtyping method, we examined 351 genomes of *C. difficile* isolates derived from a clinical cohort from Brigham and Women’s Hospital (BWH) in Boston (Fig. 2c) [55]. As they were not included in our initial database, they are ideal for testing the robustness and effectiveness of our subtype classification. All identified toxins could be accurately assigned to our reference sequences, with most (97%) aligning with 100% identity to our database, and the remainder aligning with >= 99.8% identity. Out of 351 total strains, 62 (17.7%) were toxin deficient, while 289 (82.3%) contained TcdA and/or TcdB genes (Table S4). Of these, there were 12 distinct subtype combinations, with frequencies similar to those observed in the NCBI dataset. A1/B1 strains were most common (N = 222), followed by A2/B2 (N = 24), - /B3 (N = 11), and A3/B5 (N = 10) (Fig. 2c). Therefore, our method was able to rapidly and automatically classify a large dataset of 351 clinically relevant *C. difficile* isolates, with all sequences represented in our current classification.

### Intragenic recombination drives TcdB diversification

We next focused on understanding the evolution of TcdA and TcdB variants and mechanisms for their diversification. To visualize global patterns of variation within TcdA and TcdB, we developed a haplotype coloring algorithm (https://github.com/doxeylab/haploColor) based on previous methods for genome visualization [56]. First, sequences are painted black where they matched the reference sequence (i.e., B1.1). Then, remaining positions were painted different colors where they matched selected other subtypes (Fig. 3a): blue when matching B5.1, gold when matching B4.1, and green when matching TcsL. The result of this algorithm applied to the TcdB alignment revealed a striking block-like and highly mosaic pattern of amino acid variation, which strongly indicates recombination between subtypes (Fig. 3a). B1, B5, and B6 are composed of a B1-like variation (black) pattern across their full-length sequences, while B4 and B7 are composed of a B4-like pattern (gold) across their full-length sequences. B2, B3, B8, and B9, however, possess a mosaic combination of B1-like and B4-like patterns. B3, B4, B7, and B8 share a distinct B4-like pattern of amino acid variation across their N-terminal region including the GTD and CPD domains, but when examining the DRBD, the B4-like pattern is shared by a different set of subtypes (B2, B4, B7, B10, and B11). These patterns indicate ancestral within-gene (“intragenic”) recombination events involving distinct regions of TcdB. As a statistical test of recombination, we further performed phylogenetic network analysis using SplitsTree [57]. Consistent with patterns of amino acid variation and per-domain phylogenetic analysis, network analysis revealed significant evidence of recombination within TcdB (*p* = 0; Phi test for recombination) (Fig. S4). In contrast to TcdB, TcdA (1-1874) produced homogeneous patterns of variation across each subtype (Fig. S5) and did not display evidence of recombination in network analysis (*p* = 0.186) (Fig. S4), indicating that recombination occurs frequently only in TcdB, but not in TcdA.

**Figure 3.**
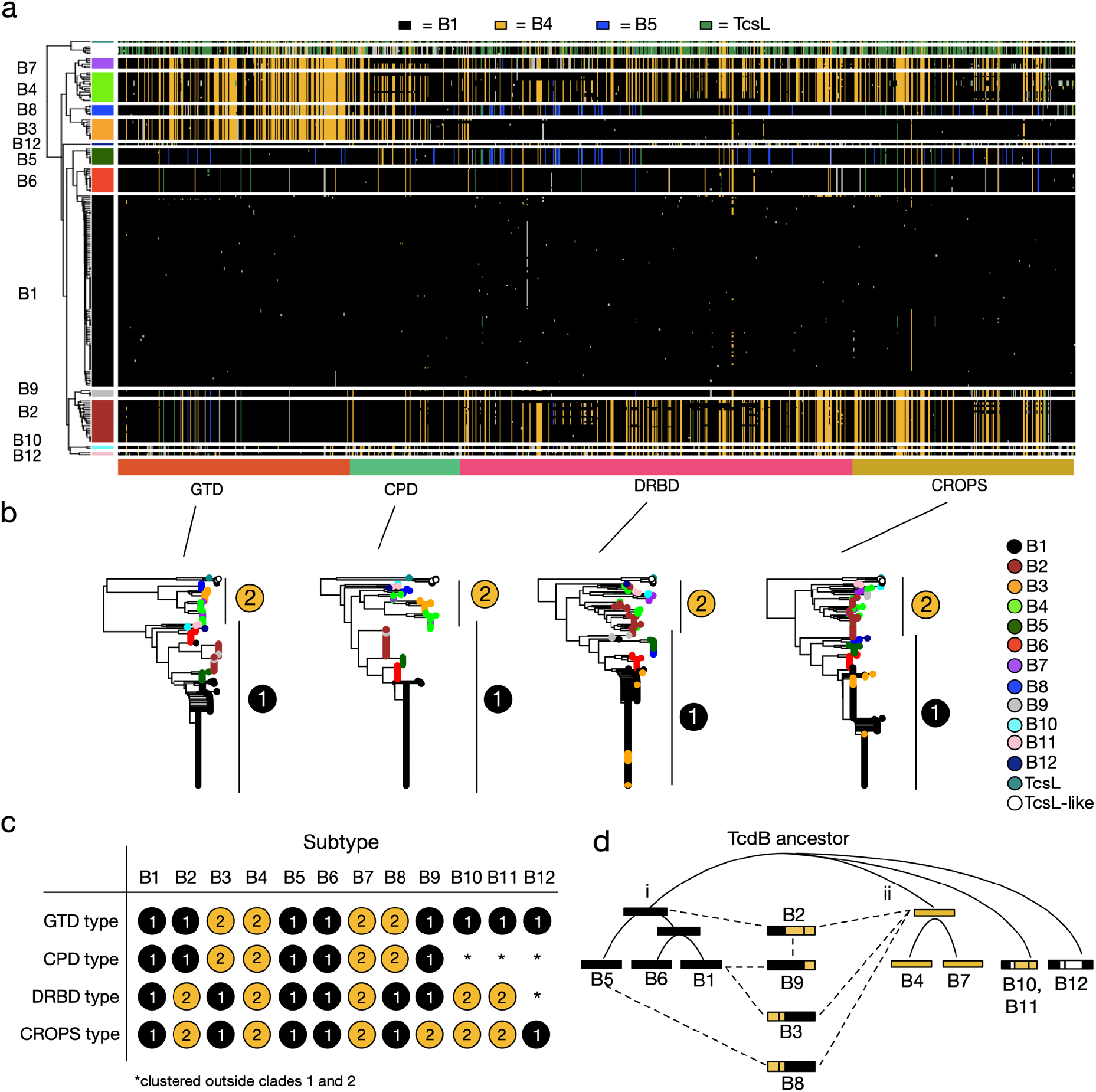
Evolutionary diversification of TcdB by intragenic recombination and domain shuffling. (**a**) Visualization of amino acid variation patterns in TcdB using a newly developed haplotype coloring algorithm (HaploColor). The visualization shows patterns of amino acid variation across the TcdB alignment. In this algorithm, the first sequence (B1.1) is assigned a distinct color, and all other sequences are colored the same color where they match this first sequence. Then, the process is repeated using a second sequence (B4.1) as the new reference, and so on. This reveals multiple colored segments indicative of common ancestry (identity by descent). Mosaic patterns are indicative of intragenic recombination. **(b)** Phylogenetic trees of TcdB based on individual domains. Each domain tree can be subdivided into two types (labeled 1 and 2), which allows each subtype to be described based on its domain composition (**c**). This reveals that TcdB subtypes are composed of domains with variable evolutionary histories, indicative of domain shuffling and intragenic recombination. (**d**) Evolutionary model depicting relationships between subtypes and putative recombination events. Here, TcdB split early into two main groups (i and ii). Subtype B2 likely originated by a recombination event fusing an ancestral type i and type ii toxin. B9 likely originated from recombination between B1 and B2, B3 from recombination between B1 and a type ii toxin, and B8 from recombination between B5 and a type ii toxin.

We further performed separate phylogenetic analyses of each domain (GTD, CPD, DRBD, and CROP) of TcdB (Fig. 3b). The phylogenetic tree of each domain produced two main groups (labeled i and ii), which correspond with the B1-like and B4-like patterns revealed in the alignment visualization (Fig. 3a). Each subtype can therefore be described as a chimeric combination of type “i” (B1-like) or type “ii” (B4-like) domains (Fig. 3c). Based on the per-domain phylogenetic relationships and recombination patterns, we formulated a potential evolutionary model for the origin of TcdB subtypes (Fig. 3d). An early TcdB ancestor split into two main groups: (i) B1, B5, and B6; and (ii) B4 and B7. Subtype B2 likely originated by a recombination event fusing an ancestral type i and type ii toxin. B9 likely originated from a recombination event between B1 and B2, B3 from a recombination event between B1 and a type ii toxin, and B8 from a recombination event between B5 and a type ii toxin. Subtypes B10-B12, which are recently identified rare variants, are early diverging lineages since they consistently outgrouped other subtypes in phylogenetic analysis (Fig. 3b), consistent with their divergent lineages among other strains (Fig. 2a).

In addition to these major ancestral recombination events, we also identified a considerable degree of “microrecombination” events involving exchange of small segments between subtypes. For example, a single TcdB sequence (B1.59) from subtype B1 has acquired an N-terminal segment that is clearly derived from subtype B2 or B9 (Fig. 3a, Figure S6). This unique TcdB gene, which appears to be the result of a spontaneous recombination event between a B1 and B2-containing strain, is derived from a newly assembled clinical isolate from a Fidaxomicin clinical trial (SRS1378602). A second similar example is B1.58 from a clinical isolate (ECDC-040, SRS1486176), which has acquired a DRBD and CROPS segment from a B2-containing strain (Figure S6). Fourteen such cases of microrecombination including these are depicted in Figure S6. TcdB in particular appears to have diversified through an extensive degree of intragenic recombination involving both large and small segments.

### Subtyping partitions TcdB into distinct functional and antigenic groups

The value of subtyping classification is to facilitate a molecular understanding of the impact of sequence variations on function and antigenicity. For instance, our sequence alignment divides the GTD into two groups: one contains B3, B4, B7, and B8; and the rest form another group (Fig. 3b, Fig. S7). Previous studies have reported two types of cell-rounding effects: TcdB1 and B2 are known to induce rounded cells with many protrusions remaining attached to cell culture plate, whereas TcdB from the strain 1470 and 8864 have been reported to cause rounded cells without protrusions, which is similar to TcsL [58]. It has been proposed that this is a result of the altered specificity of their GTD in targeting different small GTPases [32,58]. TcdB in strain 1470 is classified as B3, and the strain 8864 expresses B7, thus our classification predicts that the group containing B3/4/7/8 induces TcsL-like cell rounding phenotype. This is indeed the case for two recently reported clinical strains HSJD-312 and HMX152: both express toxins classified as B4 under our subtyping system (Table S2) and have been reported to induce TcsL-like cell rounding [59].

Another well-characterized functional motif in TcdB is its FZD-binding interface, with key residues clearly defined by the co-crystal structure [42,43,60,61]. It has been reported that B2 lacks the ability to bind FZDs due to residue variations at FZD-binding interfaces [40,44,45]. To survey whether these variations may also exist in other subtypes, we aligned the key residues across all TcdB sequences and visualized them. As shown in Fig. 4a, the FZD-binding motif is highly conserved across B1/3/5/6/8/9, while B2/4/7/10 share the same set of residue changes. Thus, B4/7/10 are predicted to lack FZD-binding capability similar to B2. B11 contains a subset of residue changes found in B2 within this region and likely also has reduced binding to FZDs. This pattern is consistent with the phylogenetic alignment of the DRBD domain, in which B1/3/5/6/8/9 form group i and B2/4/7/10/11 form group ii (Fig. 3b, 3c). Interestingly, although most B2 variants possess FZD-binding site substitutions, there are a few exceptions that contain a largely intact FZD binding site. In particular, B2.12 assembled from strain 2007223 (ERS001491) contains only a single amino acid substitution (F1597S) in this region. Examination of the alignment reveals that this is likely due to a microrecombination event that has replaced most of the FZD binding site with a B1-like segment (Fig. 4a, Fig. S6). A similar scenario occurred in a member of subtype B6, in which a B1-like segment has partially replaced this region (Fig. S6).

**Figure 4.**
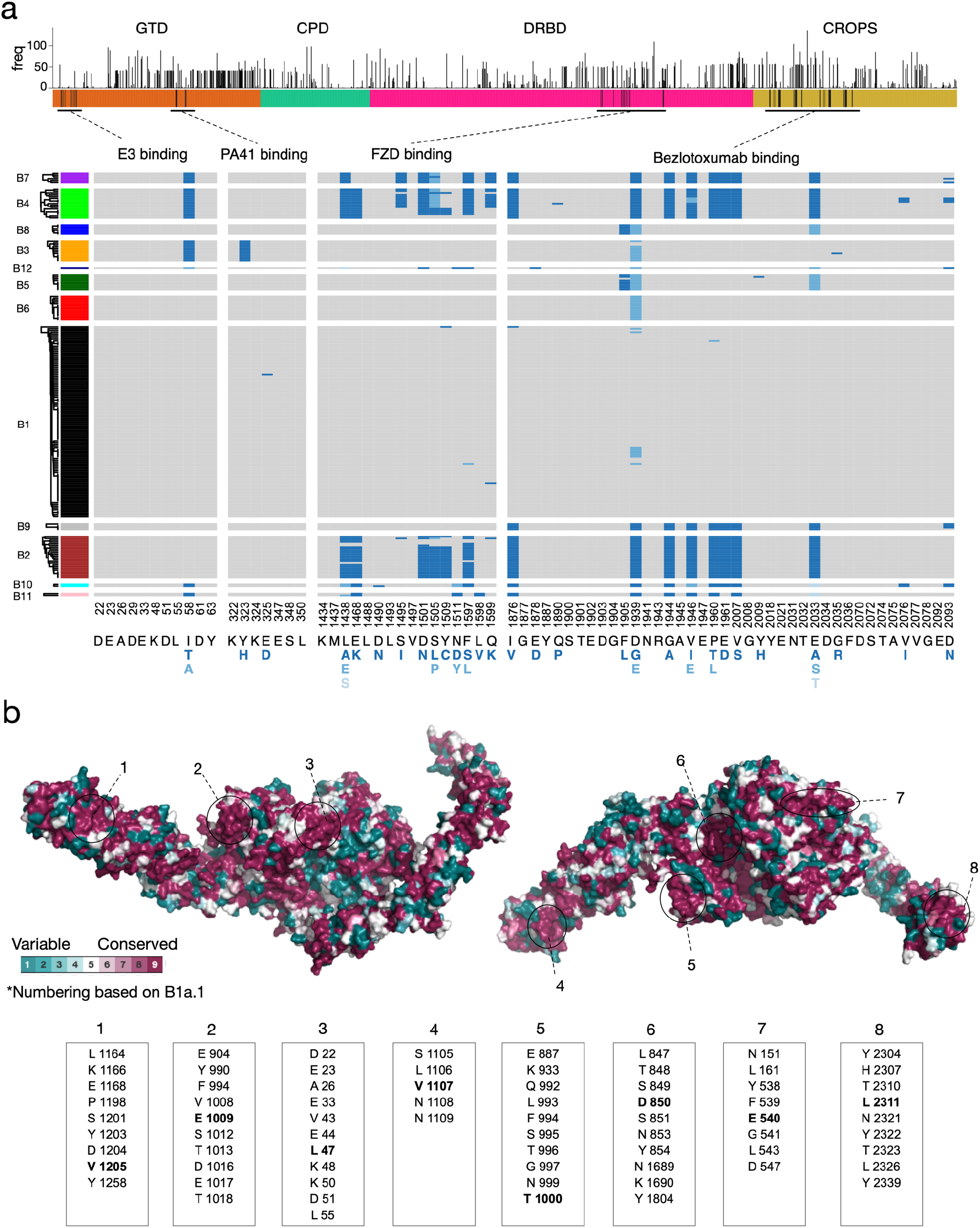
Conservation and functional variation across TcdB subtypes. (**a**) Frequency of amino acid variants across all positions of TcdB. The height of the bar indicates the number of unique TcdB sequences that contain a substitution relative to the classical TcdB1 (B1.1) sequence from strain 630 and VPI10463. Below this is a plot of amino acid variation for key functional regions including the binding sites for the frizzled receptor (FZD) and the antibodies (E3, PA41, and bezlotoxumab). The alignment is colored gray for residues that match the common amino acid found in B1.1, and variants are colored blue (darkest blue = most common variant). E3 and PA41 binding sites are highly conserved, whereas FZD and bezlotoxumab binding sites are highly variable. FZD and bezlotoxumab variants also co-occur with each other. (**b**) Evolutionary conservation mapped to the protein structure of full length TcdB based on PDB 6OQ5 [65]. Eight highly conserved surface patches are indicated. Center residues within each surface patch are indicated in bold font.

Sequence variations between subtypes could also have a drastic impact on the efficacy of therapeutic antibodies and vaccines. Bezlotoxumab from Merck is the only monoclonal antibody against TcdB that has so far been approved by the FDA and is currently used to reduce the recurrence of CDI [62]. This antibody was generated using fragments of TcdB1 as antigens and its epitope sites (located at the N-terminal of CROPs) have been established through crystallography [63]. We thus aligned all key residues within its epitope across all TcdB sequences, which revealed extensive residue changes largely conserved in B2/4/7/9/10/11 (Fig. 4a). This is consistent with our alignment of the CROPs domain that group B2/4/7/9/10/11 together (Fig. 3b, 3c). It has been shown that bezlotoxumab exhibited as low as ~200-fold reduction in neutralization efficacy against TcdB from several RT027 strains, which likely express B2, compared with its efficacy against B1 from VPI10463 [41,64]. It also showed a similarly low efficacy against a strain 8864, which expresses B7. These results indicate that bezlotoxumab does not have good efficacy against CDI caused by strains that express B2/4/7/9. Furthermore, there are also a few amino acid changes within the epitope region in B5/B8, and it has been shown that bezlotoxumab has ~60-fold reduction in efficacy against the TcdB from a RT078 strain [41], which likely express B5 (Table S1). These results clearly indicate that subtype classification of toxins will be able to guide the use of bezlotoxumab in the clinic.

In addition to bezlotoxumab, we also examined another monoclonal antibody PA41, which is under development [41], and a single-domain antibody (also known as VHH or nanobody) E3 [65]. The epitopes for both have been well established through co-crystal structures [65,66]. Both recognize the GTD domain, with E3 recognizing the N-terminus of TcdB (Fig. 4a). The epitope site for PA41 is highly conserved across most subtypes except a single residue change (Y323H) in B3. This is consistent with the previous finding that PA41 can potently neutralize TcdB from many different strains except RT017 strains, which express B3 [41]. The epitope site for E3 is conserved in most subtypes except a single residue change (I58T or A) in B3/4/7/8/10/11/12, and the impact of this single residue change remains to be examined experimentally.

Finally, we mapped evolutionary conservation across all available TcdB sequences onto the recently reported crystal structure of TcdB [65] (Fig. 4b). Relative to the reference TcdB1 sequence, amino acid variants are common across the full-length TcdB sequence and occur throughout each domain (Fig. 4a) but some regions (e.g., N-terminus of the GTD, C-terminus of CROPS domain, segments of the pore-forming region of the DRBD and C-terminus of the CROPS domain) were highly conserved. Based on structure, we identified eight conserved surface patches containing universally conserved residues which represent potential key therapeutic targets for developing broad-spectrum diagnostics, antibodies, and vaccines (Fig. 4b).

### Diff-base: a central hub for storing and analyzing TcdA and TcdB sequences

To address the needs of the research and clinical community in understanding toxin subtyping and variations, we developed an online open database freely accessible at *<u>diffbase.uwaterloo.ca</u>*. DiffBase stores all unique TcdA and TcdB sequences identified to date from the NCBI and SRA and organizes sequences into our subtype classification scheme. Different subtypes and individual sequences can be explored and visualized in reference trees, with additional information such as source strains, and links to other resources (Fig. S8). In addition, users can query their own TcdA or TcdB sequences against the database using a built-in BLAST interface, which will report the top matching sequences in the database and provide toxin classifications and other related information. To keep up with new sequences and information concerning TcdA and TcdB, DiffBase facilitates community feedback and allows users to submit new information to be added to the next iteration of the database.

## Discussion

Here we created the largest database to date capturing available TcdA and TcdB sequence diversity. This up-to-date collection includes genes from sequenced *C. difficile* isolates in GenBank, as well as thousands of genomes that were assembled, annotated and analyzed from the NCBI short-read archive. We clustered TcdA and TcdB variants into phylogenetic subtypes, which provided a robust classification that is both congruent with *C. difficile* genome phylogeny as well as variation in functional and therapeutically relevant amino acids including TcdB regions targeted by existing monoclonal antibodies. Our analysis revealed that TcdB undergoes extensive homologous recombination, and its potential evolutionary history is proposed based on recombination among various subtypes. Finally, our analysis revealed mapped eight conserved patches across the TcdB structure, which will facilitate future studies that aim to develop “universal” *C. difficile* therapeutics that broadly target all TcdB subtypes.

In general, there is some agreement between previously defined toxinotypes and our toxin subtypes, but subtyping provides additional information as it is able to describe TcdA and TcdB separately. For example, toxinotype 0 associates largely with the A1/B1 subtype, toxinotype III associates largely with A2/B2, toxinotype IV with A3/B5, toxinotype VIII associates with -/B3, toxinotype IX associates largely with A2/B4, toxinotype X associates with -/B7, and so on (Table S1). However, subtype A3/B5 associated with toxinotypes V, VI, VII, XVI, XXVIII, all of which are found in clade 5 strains. Moving forward, with improved abilities to perform genome sequencing of clinical isolates, it will be increasingly possible to classify strains based on their genome-wide phylogenetic relationships as well as their toxin subtypes.

In comparison to TcdA, our analysis identified a much greater degree of sequence variation within TcdB and a larger number of subtypes. Given that we see evidence for extreme recombination in TcdB but not TcdA, it is possible that there is a greater selective pressure for positive selection and diversification of TcdB. We speculate that intragenic recombination of TcdB may drive antigenic diversification, whereas in TcdA this process may be driven by truncation and variation of its C-terminal CROPS region. The CROPs domain showed similarity with carbohydrate-binding proteins and may contribute to toxin attachment to cells by binding to carbohydrate moieties (27, 42, 44). The CROPs domain may also act as a chaperone that protects other domains (45). Possibly due to its repetitive nature, the CROPs domain is often the region that induces strong immune responses. It remains to be determined whether frequent recombinations/changes in TcdA-CROPs may alter its function and/or antigenicity.

These findings further suggest that TcdB may play a central role in *C. difficile* pathogenesis, which is consistent with previous findings that TcdA-/TcdB+ mutant *C. difficile* strains are fully virulent, whereas TcdA+/TcdB- strains are attenuated in multiple mouse models [26,67]. It has also been suggested that TcdB is the primary factor for inducing the host immune and inflammatory responses in mouse models [26]. The key role of TcdB in CDI is further confirmed by the findings that an antibody that neutralizes TcdB (bezlotoxumab), but not another one that neutralizes TcdA (actoxumab, Merck), conferred protection against CDI in gnotobiotic piglets [68] and reduced CDI recurrence in humans [62,69] and it is also consistent with the fact that many clinical isolates only express TcdB [70]. An exception to a dominant role for TcdB is the very rare TcdA+ TcdB-strain. It is noteworthy that one such strain identified in GenBank contains the single most divergent TcdA sequence (subtype A7) [53], which may have diverged to acquire a pathogenic functionality without requiring TcdB.

For such recombination events to have occurred in TcdB, it is possible that phylogenetically distinct *C. difficile* strains containing different toxin subtypes coexisted within the same host individuals, exchanged genetic material and recombined to produce new recombinant forms. Coinfection with different *C. difficile* ribotypes has been recently reported in a clinical case study [71]. Theoretically, co-infection does not need to occur frequently to promote recombination. A single individual containing two or more *C. difficile* strains may be sufficient to promote recombination, generating hybrid toxins with different regions derived from different subtypes. The new recombinant strain can then increase in frequency through genetic drift and/or selection, as well as through transmission to other individuals. Our analysis suggests that this process has not only occurred frequently in the past as a mechanism by which different subtypes originated, but that it may be a frequent and ongoing process in new clinical isolates (e.g., B1.59 from SRS1378602). Consideration of intragenic recombination and how it may shape TcdB function and toxicity will be important in efforts to understand the emergence of new *C. difficile* hypervirulent strains and develop targeted therapeutic interventions.

Recombination offers considerable adaptive benefits to proteins by facilitating rapid mutation of a sequence by exchange of entire segments as opposed to the relatively slower process of single point mutations. In this way, proteins can diversify by shuffling a few basic building blocks such as protein domains. In pathogens, recombination plays a major role in pathoadaptive evolution by facilitating rapid “switching” of virulence factors and antigenic proteins [72,73]. Antigenic recombination can promote the sudden avoidance of immune recognition (antigenic escape), which has been demonstrated for the *C. difficile* S-layer gene [74]. In the case of TcdB, intragenic recombination may generate new hybrid toxins composed of different domains types and functions. In theory, recombination could also generate resistance to therapeutics by replacing entire binding interfaces with compatible regions from other toxins that possess drug-resistant mutations. Recombination-mediated domain shuffling not only describes TcdB sequence patterns and phylogenetic relationships, but also provides an explanation for important functional differences between TcdB variants. For example, the exchange of a B4-like GTD between subtypes B3, B4, B7 and B8, correlates with the TcsL-like clumping and rounding phenotype. Also intriguing are the many microrecombination events that have occurred in the DRDB region which overlap with FZD-binding site. For example, likely due to partial homologous recombination with a B1-like toxin, one B2 variant (B2.12, strain 2007223 from ERS001491) contains an intact FZD-binding interface with only a single amino acid substitution (F1597S). This suggests that intragenic recombination in TcdB may promote rapid evolutionary switching between receptor-binding activities or affinities.

Given the extent of TcdB diversification and its primary role in virulence, it is critically important to identify conserved regions that can be targeted for therapeutic and diagnostic applications. Sequence conservation mapped to protein structure also revealed at least 8 distinct surface patches containing a high density of universally conserved residues across all TcdB subtypes, which represent promising regions for the development of inhibitors. Importantly, the binding site for the antibody therapeutic bezlotoxumab, which is commonly used to treat *C. difficile* infections, was not among these and instead displayed considerable variation across TcdB subtypes with B2, B4, B7, B9, B10, B11, in particular displaying 7-8 likely destabilizing substitutions. Although the common B1 subtype of TcdB is largely conserved across this region, based on analysis, it is possible that intragenic recombination with other strains (e.g., a B2-containing strain) could generate spontaneous resistance to bezlotoxumab by replacing this region with a B2-like segment. Future efforts to target highly conserved clusters of surface-exposed residues on the TcdB structure may yield promising candidates for therapeutic or vaccine development.

Finally, based on sequence-based classification of *tcdA* and *tcdB* genes, we propose a revised scheme for naming these genes in future studies. In this scheme, a newly identified TcdA or TcdB sequence may be aligned to our reference database and named based on the top hit according to sequence identity. In order to enable automated subtyping of new *tcdA* and *tcdB* genes and facilitate community collaboration and data sharing, we have developed a freely available online database (DiffBase) for use by the *C. difficile* clinical and research community. In the future, clinicians will be able to query toxin sequences from clinical isolates and immediately determine the toxin subtype, which will help them decide on therapeutic strategies. For instance, among the 351 clinical cases in the BWH dataset, there are 34 cases expressing B2/B4/B7/B9, for which treatment of bezlotoxumab would not be effective. Therefore, toxin subtyping will guide proper choices of clinical treatment in consideration of toxin variations and allow researchers to monitor the ongoing evolution and diversification of *C. difficile*.

Note: Related work was published during the preparation of this manuscript that performed a subtype analysis of TcdB [50]. The analysis was limited to TcdB only and also included a smaller set (3,269) of genomes, resulting in fewer (8) TcdB subtypes. Their classification is consistent with our analysis, but does not capture the full diversity of available sequences and subdivide sequences according to a comprehensive analysis of recombination patterns. Furthermore, no online database was provided to facilitate automated subtyping analysis. Our work is also unique through its model of toxin evolution, analysis of conserved and variable regions and their impact on antigenicity, discovery that the current therapeutic antibody is only effective on a subset of toxins, and its analysis of a validated clinical dataset from Brigham and Women’s Hospital.

## Methods

### Dataset construction

#### Assembly of 6492 C. difficile genomes from the NCBI short read archive

A set of *Clostridiodes difficile* sequencing runs was retrieved from the NCBI short read archive (SRA) by text query for *“Clostridioides difficile”* on June 20th, 2019. Metagenomic samples were omitted, leaving only genomic samples to reduce the chance of contamination from other bacterial species. Sequencing runs were downloaded using the fasterq-dump module of the SRA toolkit. To account for multiple library preparation methods and adapters, the fastp tool [75] was used to perform adapter trimming and quality control of the sequencing reads. For each quality-controlled set of reads, SPAdes version 3.12 [76] was used for genomes assembly, with *C. difficile* str. 630 as a conservative reference and the --untrusted-contigs and --careful options. Each assembly was automatically annotated using the Prokka pipeline [77] with a minimum contig length of 200. In order to verify the identity of the assembled genomes as strains of *C. difficile*, the predicted genes from Prokka were taxonomically annotated using Centrifuge [78] against their pre-compiled index of bacterial, archaeal, viral, and human genomes. Only samples that were clearly identifiable as strains of *C. difficile* were kept.

To identify the *tcdA* and *tcdB* genes from all strains, the phmmer tool was used to search for matches to TcdA (uniprot accession # Q189K5_CLOD6) and TcdB (uinprot accession # Q189K3_CLOD6) as queries. In order to distinguish true sequence variants from poorly assembled, low-quality, or chimeric variants, only hits that clearly represented well-assembled toxin sequences (that is, yielding a protein equal to or greater than 1,800 amino acids in length) were retained. Sequences with apparent N- or C-terminal truncations representing less than 1% of the total assembled data set were also removed. In total, the final re-assembled set of redundant TcdA and TcdB sequences consisted of 3,542 and 5,994 sequences, respectively. Redundancy in each of these sets was removed by clustering with CD-HIT version 4.6 [79] at 100% identity. Non-redundant sets were aligned using the L-INS-i algorithm of the MAFFT package [80].

#### TcdA and TcdB sequences from the NCBI GenBank database and manually curated set

GenBank homologs of TcdA and TcdB were also identified via a BLAST search of the NCBI non-redundant database on Feb 8, 2020. TcdB and TcdA sequences from *C. difficile* strain 630 were used as queries. Homologs were filtered to those with *E*-value < 1e-10, 70% identity and query alignment coverage, which removed partial sequences. In addition, we manually curated 63 reference *C. difficile* strains collected from previous studies [12,46]. For these 63 genomes, we manually identified corresponding strains within the NCBI or SRA database and identified *tcdA* and *tcdB* genes based on pre-computed genome annotations or through similarity searches. Fourteen genomes could not be associated with *tcdA* and *tcdB* genes in the NCBI; for these cases, raw genomic reads were retrieved from European Nucleotide Archive (ENA) and were assembled using SPAdes as described earlier.

#### Construction of TcdA and TcdB alignments

A combined dataset of TcdA and TcdB homologs was created by pooling SRA-derived, NCBI-nr derived, and the manually curated set of sequences. The combined set of sequences were aligned using MUSCLE [81] with default parameters as implemented in Seaview [82]. Due to significant length variation at the C-terminus of TcdA alignment, only the CROP-less core region (1-1874) of the alignment was kept for subsequent analysis; while the entire TcdB alignment (1-2366) was used. Redundant sequences (100% identity) were removed as well as sequences annotated as partial that contained truncations in the alignment.

### Sequence clustering and analysis

TcdA and TcdB alignments were then processed separately using an analysis pipeline implemented within R. For each case, the multiple sequence alignment was converted to a distance matrix using the dist.alignment() function from the seqinr package [83]. Average linkage hierarchical clustering was performed using the hclust() function. Pairwise sequence similarities were mapped onto the clustering tree and visualized using the ComplexHeatmap package [84] and clustering threshold were chosen to generate subtypes with strong internal (within-cluster) and lower external (between-cluster) similarities based on visual analysis and quantitative analysis of percentage identity distributions.

For analysis of amino acid variation, we converted the alignments into data matrices using the alignment2matrix() function from the BALCONY R package [85]. We then identified all variant residues across all alignment positions relative to the sequences of TcdA and TcdB from strain 630 as a reference. Residues implicated in frizzled binding [42], bezlotoxumab binding [63], PA41 binding [66], and E3 binding [65] were then analyzed in terms of their variation across subtypes. E3 binding residues were identified by analysis of PDB structure 6OQ5 [65], by selecting atoms in chain A (TcdB) within a 4 Å distance of chain E (E3) using PyMol’s distance algebra functions. The ComplexHeatmap R package was used for data visualization.

Analysis of repeats in TcdA was done using InterproScan as part of the InterPro 80.0 database [86]. The number of detected matches to ProSite’s cell wall-binding repeat profile (PS51170) was counted in the A1.1 reference sequence (UniProt P16154, 2710 aa), the longest (3070 aa) and shortest (1889 aa) variants of TcdA in our database.

### Structural analysis

To map sequence conservation on to the structure of TcdB, we used the ConSurf server [87] with the TcdB alignment as input and the recently determined crystal structure of full-length TcdB (PDB ID 6OQ5) [65] as the template. Default parameters (neighbor-joining with ML distance and Bayesian calculation of conservation scores) were used. Structural visualization was done using PyMol version 2.3.4, using the recommended script (https://consurf.tau.ac.il/pyMOL/consurf_new.py) with insufficient data hidden from the image.

### Construction and toxin subtyping of C. difficile genome phylogeny

We retrieved 2,118 assemblies for 1,934 representative *C. difficile* genomes (https://www.ncbi.nlm.nih.gov/genome/tree/535) from the NCBI. The snippy pipeline (https://github.com/tseemann/snippy) was used to map all genomes to the reference (strain 630, GCA_000003215). For phylogenetic reconstruction, we analyzed 14,194 SNPs across 88 conserved marker genes (those present in *C. difficile*) derived from the PhyEco Firmicutes dataset [88]. A phylogeny was reconstructed using RAxML with the GTR+GAMMA model [89]. All TcdA and TcdB homologs from the NCBI were then subtyped by BLAST against our database of labeled toxin subtype sequences, using only the conserved portion (region 1-1874) of the TcdA alignment, and the full 1-2366 regions from the TcdB alignments. An assignment script written in Perl was used to parse BLAST output files and assign subtypes. The subtype “X” associated with the best matching reference sequence (highest sequence identity) was assigned if the alignment coverage exceeded 90% and labeled as complete; otherwise, it was labeled as a partial sequence.

### SplitsTree analysis of recombination

TcdA and TcdB alignments were analyzed by SplitsTree version 4.0 [57]. A NeighborNet tree visualization was produced using protein maximum-likelihood distances according to the WAG model of evolution. The Phi test for recombination was performed as implemented in SplitsTree which selected a window size of 100 for TcdA with k = 3 and a window size of TcdB with k = 21.

### Haplotype visualization

For visualization of recombinant blocks and haplotype structure within TcdA and TcdB protein alignments, we developed a modified algorithm based on a previous method from Wang et al. [56] for comparative genomic visualization. An implementation of this method in the R programming language is available at https://github.com/doxeylab/haploColor. The algorithms works as follows:

1. Assign first sequence as reference.
2. Assign all residues of reference a new color *C*.
3. Assign positions in other sequences that match the reference, the same color *C*.
4. Identify sequence most dissimilar to the current reference across unassigned positions, and assign it as the new reference.
5. Repeat steps 2-3 for a defined number of iterations or until all sequences are completely colored.

The algorithm was applied directly to the TcdA and TcdB alignments and run for both 4 and 16 iterations (TcdB) and 16 iterations (TcdA).

### Development of the DiffBase web-server

The DiffBase web server was developed as an R shiny() application. Contained within DiffBase is an implementation of BLAST+. Individual sequences can be submitted to the server, where the blastp program is run to find matches from within the entirety of the server sequence repositories. An *E*-value cutoff of 1e-10 is used to filter hits, and the results are sorted by percent identity between query and target sequences. Toxin groups can also be viewed in a phylogenetic tree visualized using ggtree R package [90]. Metadata about group members was obtained from the NCBI Identical Protein Group (IPG) database. The source code is freely available at https://github.com/doxeylab/diffBase.

### Data Availability

Our open source online database is available at: https://diffbase.uwaterloo.ca and https://github.com/doxeylab/diffbase.

All source code for analyses is available at: https://github.com/doxeylab/diffBaseAnalyses

## Acknowledgments

A.C.D. acknowledges funding from the Natural Sciences and Engineering Research Council of Canada (NSERC Discovery Grant, RGPIN-2019-04266; Discovery Accelerator Supplement, RGPAS-2019-00004), and from the Government of Ontario (Early Researcher Award). A.C.D. holds a University Research Chair at the University of Waterloo.

M.D. acknowledges support from National Institute of Health (NIH) (R01NS080833, R01AI132387, and R01AI139087), the NIH-funded Harvard Digestive Disease Center (P30DK034854), and Boston Children’s Hospital Intellectual and Developmental Disabilities Research Center (P30HD18655). M.D. holds the Investigator in the Pathogenesis of Infectious Disease award from the Burroughs Wellcome Fund.

J. W. acknowledges support from the Intramural Research Program of the National Library of Medicine, NIH. L.B. acknowledges support from NIH (P30 DK034854), Hatch Family Foundation, and Brigham and Women’s Hospital Precision Medicine Institute.

M.J.M. gratefully acknowledges funding from the Japan Society for the Promotion of Science as a JSPS International Research Fellow (Luscombe Unit, Okinawa Institute of Science and Technology Graduate University).

## Supplementary Data

**Figure S1.**
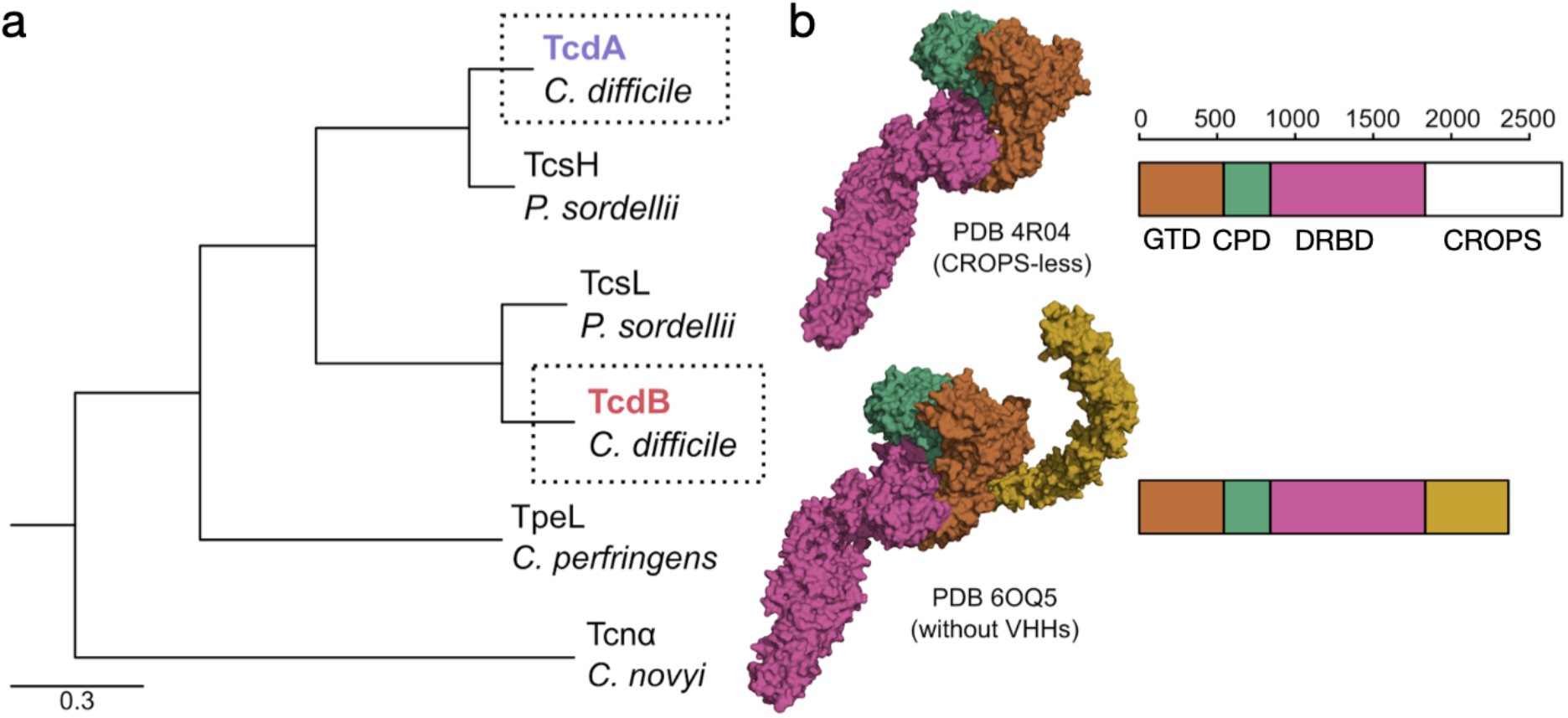
Phylogenetic and structural overview of the TcdA and TcdB protein family. (a) The TcdA family forms a monophyletic clade with TcsH from *Paeniclostridium sordellii* as a sister phylogenetic lineage. Similarly, the the TcdB family forms a monophyletic clade with TcsL from *Paeniclostridium sordellii* as a sister phylogenetic lineage. This implies a scenario whereby TcdA and TcdB evolved by an ancestral gene duplication that predates the speciation event leading to divergence of *C. difficile* and *P. sordellii*. (b) Representative crystal structures and domain architectures are shown for TcdA (above) and TcdB (below). The structure of TcdA lacks the CROPS domain and is derived from PDB ID 4F04. The full-length structure of TcdB is based on PDB ID (6OQ5), and was modified to remove bound antibodies. Domain definitions were derived from Aktories et al.

**Figure S2.**
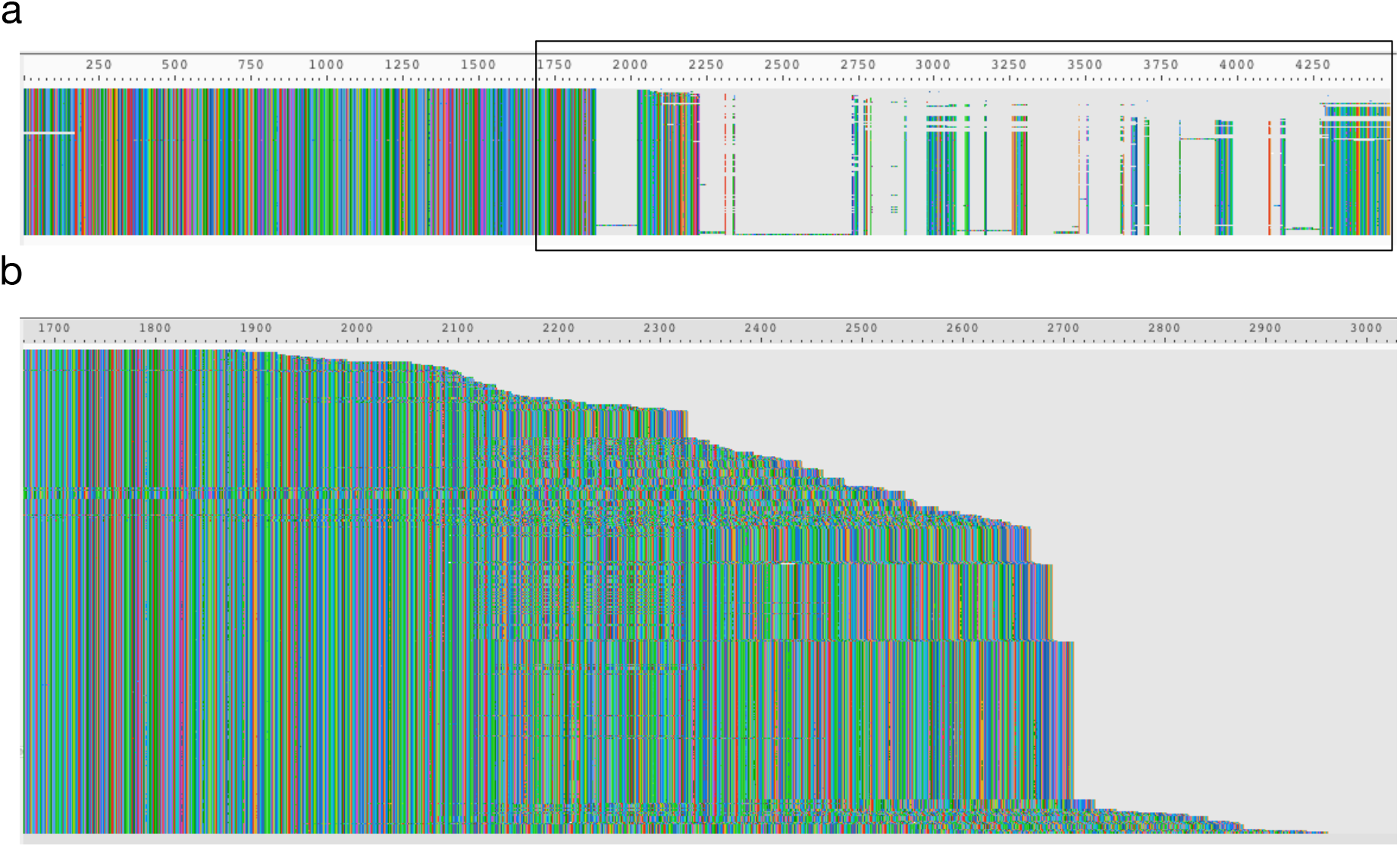
Alignment of TcdA sequences derived from GenBank and the NCBI short read archive, illustrating considerable variation in the length of the C-terminal CROPS region. (a) Complete alignment of 480 unique TcdA sequences. (b) Visualization of unaligned sequences to display C-terminal length variation following residue ~900.

**Figure S3.**
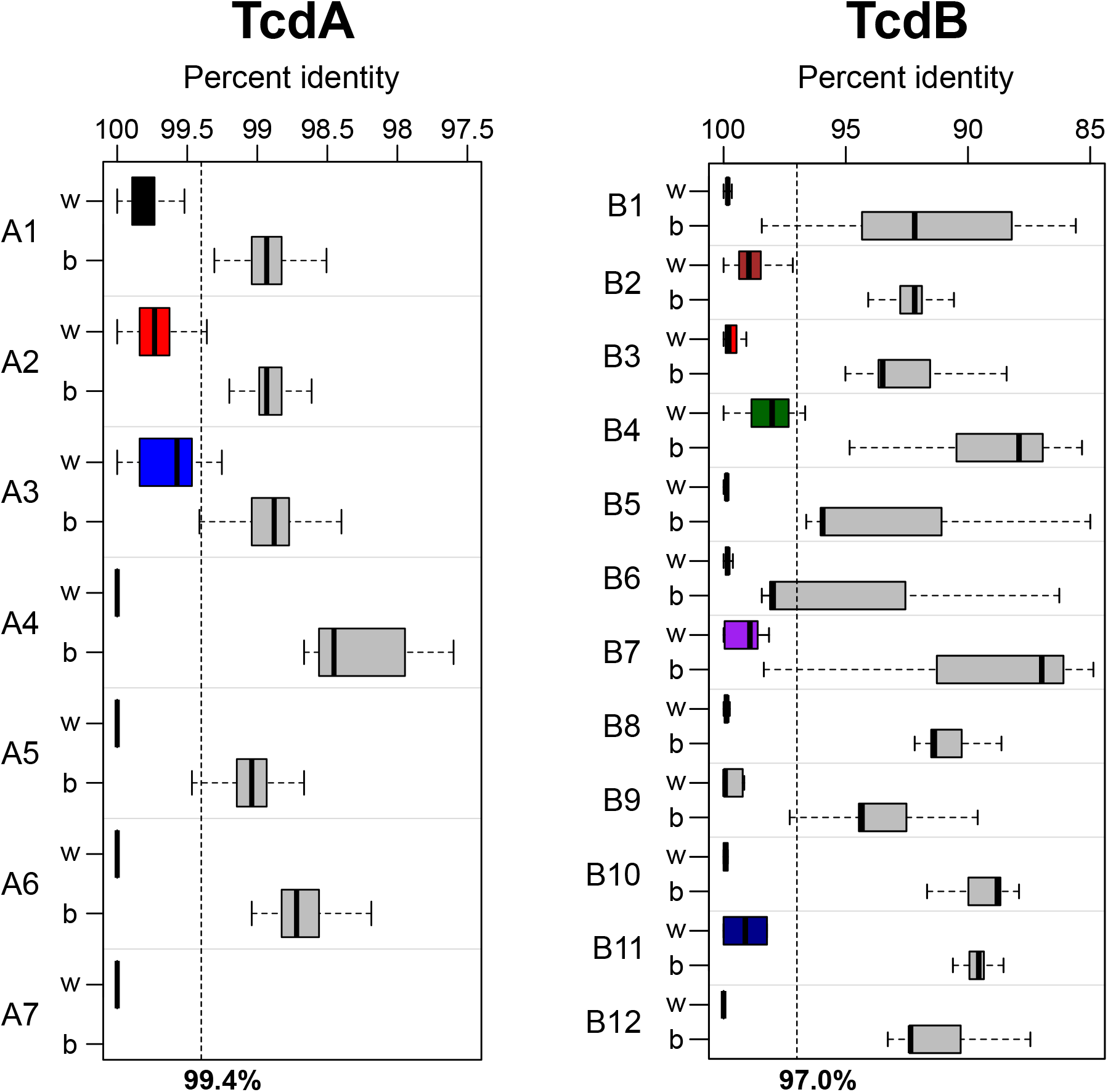
Analysis of sequence similarities within and between subtypes of TcdA and TcdB. Pairwise sequence identities were calculated between all TcdA and TcdB sequences. The % identity distributions are plotted for sequences within (“w”) the same subtype versus between (“b”) subtypes for TcdA (left) and TcdB (right). As expected, the % identities are much higher within than between subtypes. For TcdA, a % identity threshold of 99.4 effectively distinguishes sequences within the same subtype, whereas for TcdB, a threshold of 97% effectively distinguishes sequences within the same subtype.

**Figure S4.**
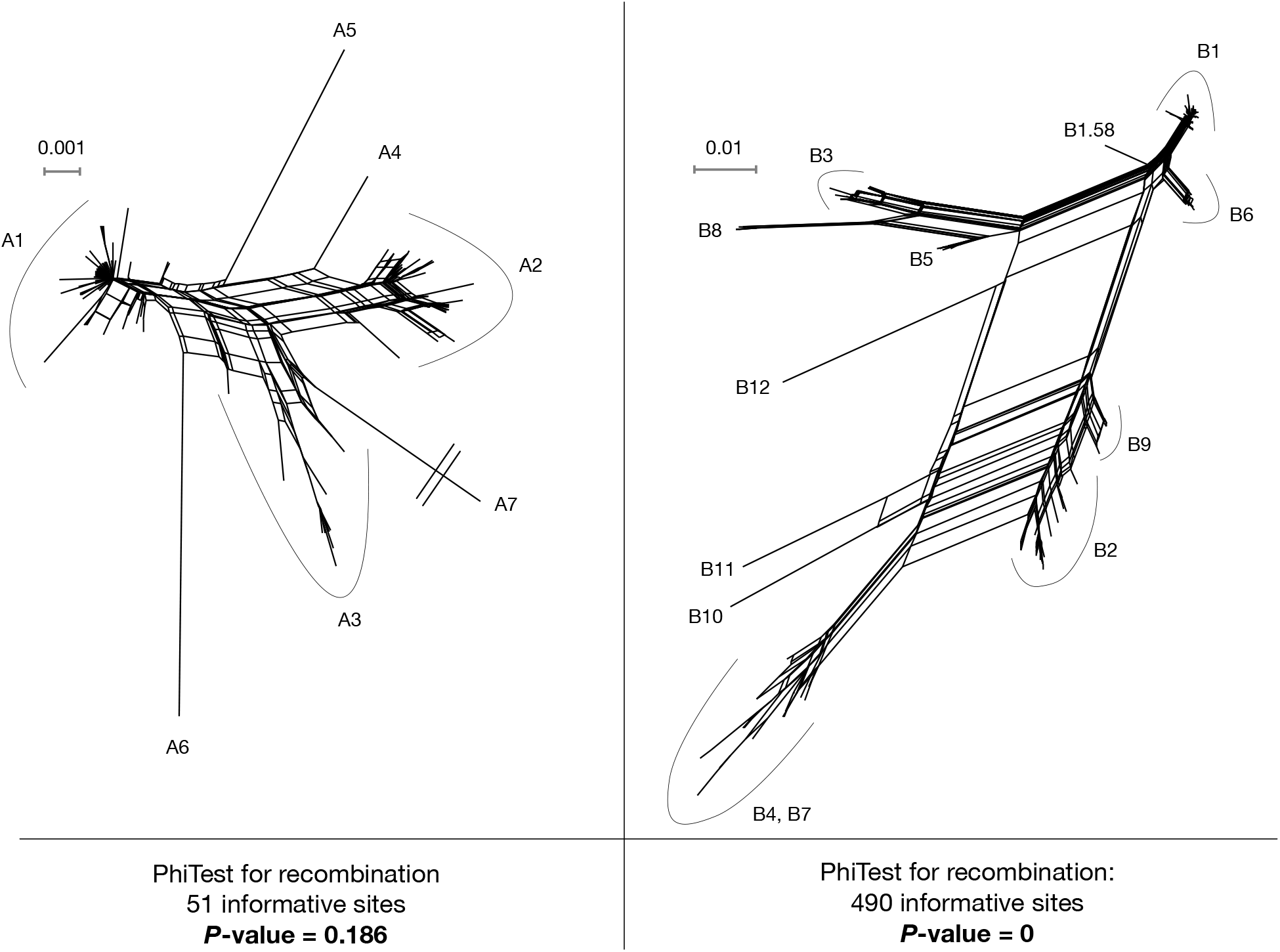
SplitsTree analysis of TcdA and TcdB and statistical detection of recombination. Split networks of TcdA and TcdB were generated using the SplitsTree software. Parallel edges suggest the existence of sites that are not compatible with a perfect monophyletic tree, which can result from recombination. An extremely long branch (A7) has been truncated in order to permit visualization.

**Figure S5.**
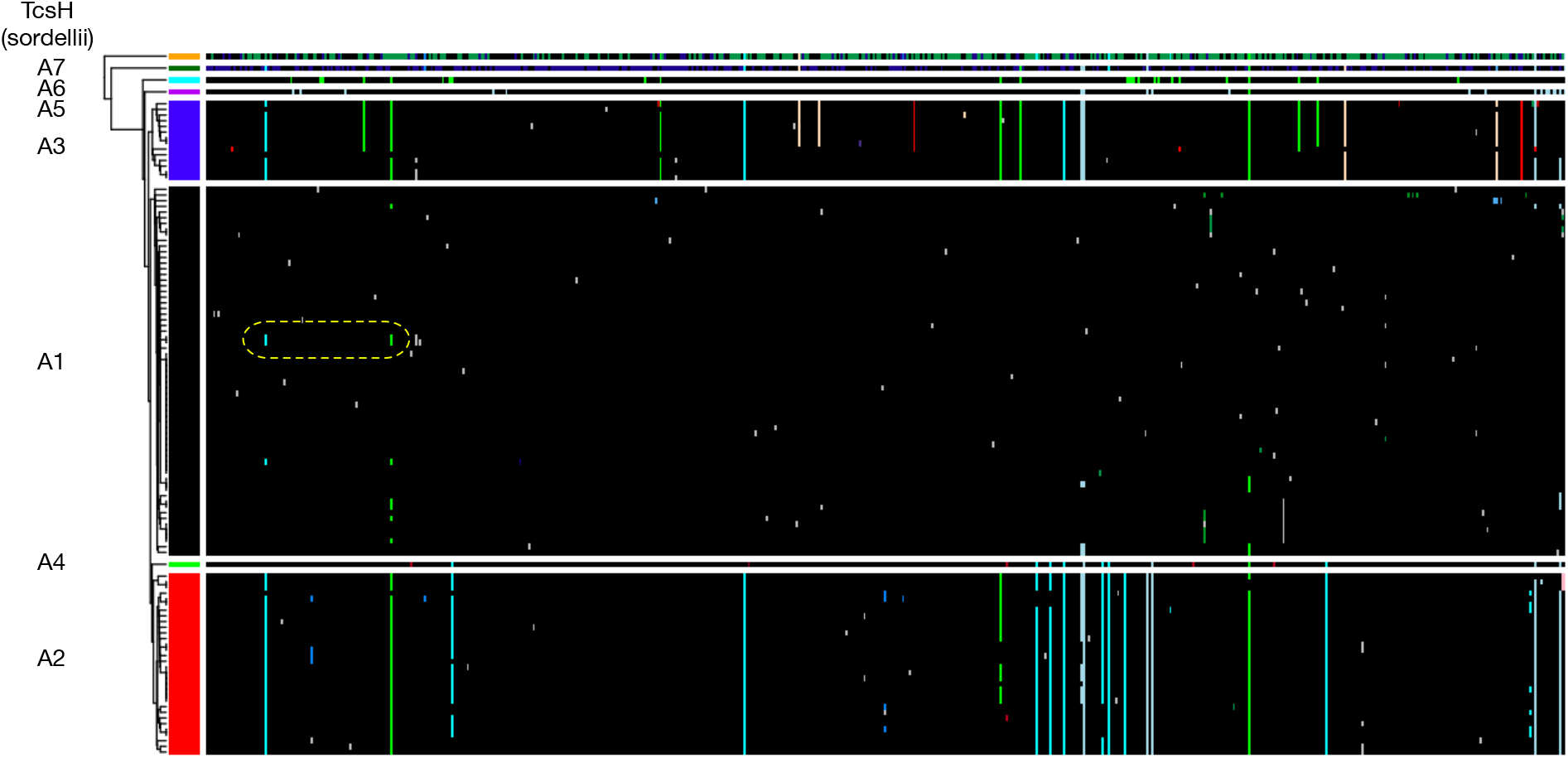
Haplotype analysis of the CROP-less TcdA alignment. Visualization and analysis of amino acid variation patterns was performed using the HaploColor algorithm (https://github.com/doxeylab/haploColor), which was run for 16 iterations. Patterns of amino acid variation within each subtype are highly homogeneous, and thus a lack of evidence for recombination. One potential exception is highlighted in yellow, involving two amino acid variants that occur within subtype A1 that are lacking in most other A1 sequences but present in subtypes A2 and A3. However, this pattern may also be due to ancestral variation rather than recombination. Overall, compared to TcdB, the TcdA displays considerably less sequence variation and lacks the mosaic patterns that would result from recombination.

**Figure S6.**
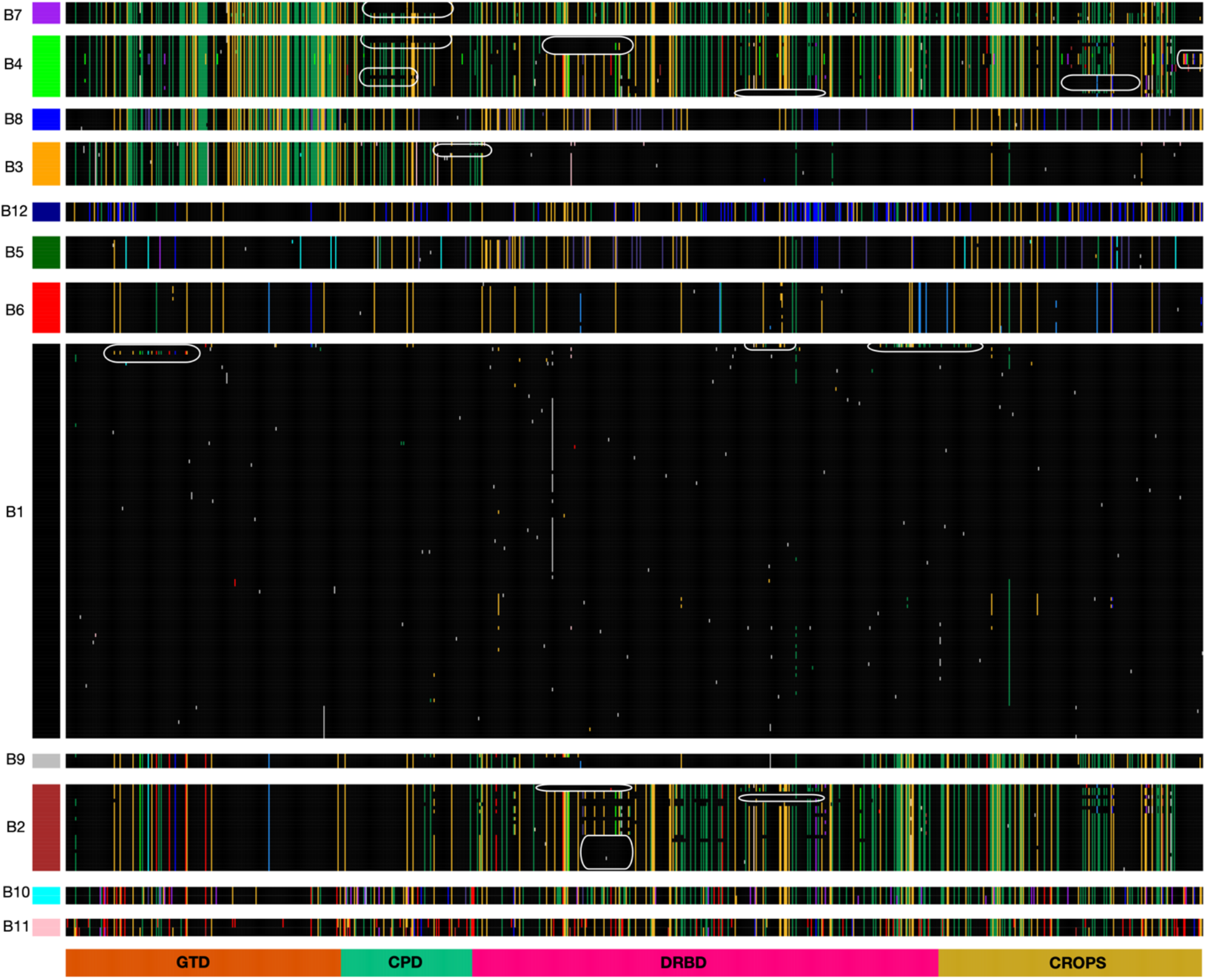
Visualization of amino acid variation patterns in TcdB highlighting putative microrecombination events. The TcdB multiple sequence alignment was colored using the HaploColor algorithm (https://github.com/doxeylab/haploColor), which was run for 16 iterations. Fourteen example segments containing amino acid variants that are unexpected for their subtype are shown by white ovals. These represent putative between-subtype microrecombination events. These fourteen are not a complete list as many more can be seen visually.

**Figure S7.**
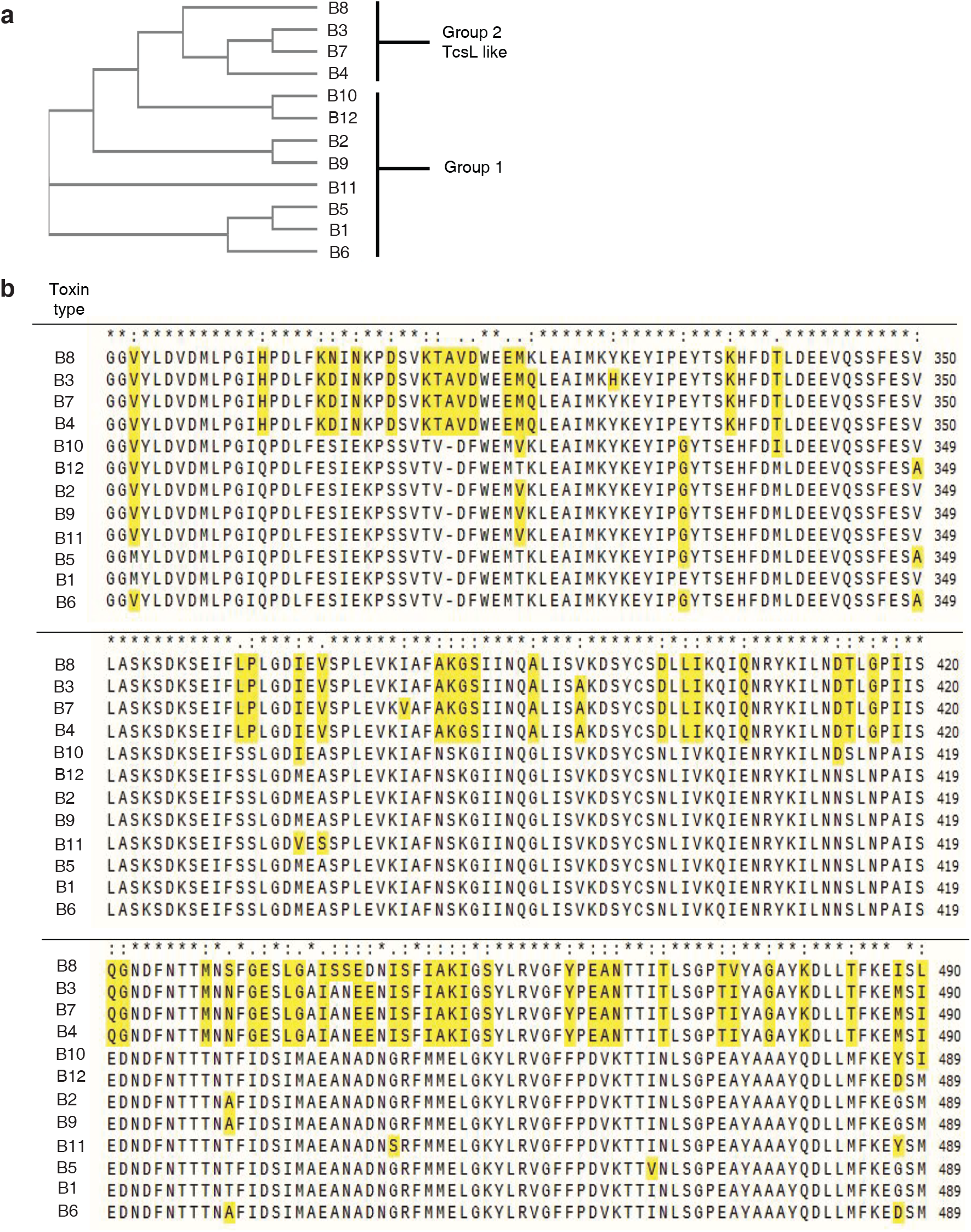
a) Phylogenetic tree of the GTD domains. b) alignment of region 280-490. According to the tree and the alignment, and the functions, GTD domains could be classed into two groups, one group is TcsL-like which gives vero cells rounding and clumping phenotypes (strain 1470 and 8846); the other group is the classical TcdB-like group which only give the rounding phenotype.

**Figure S8.**
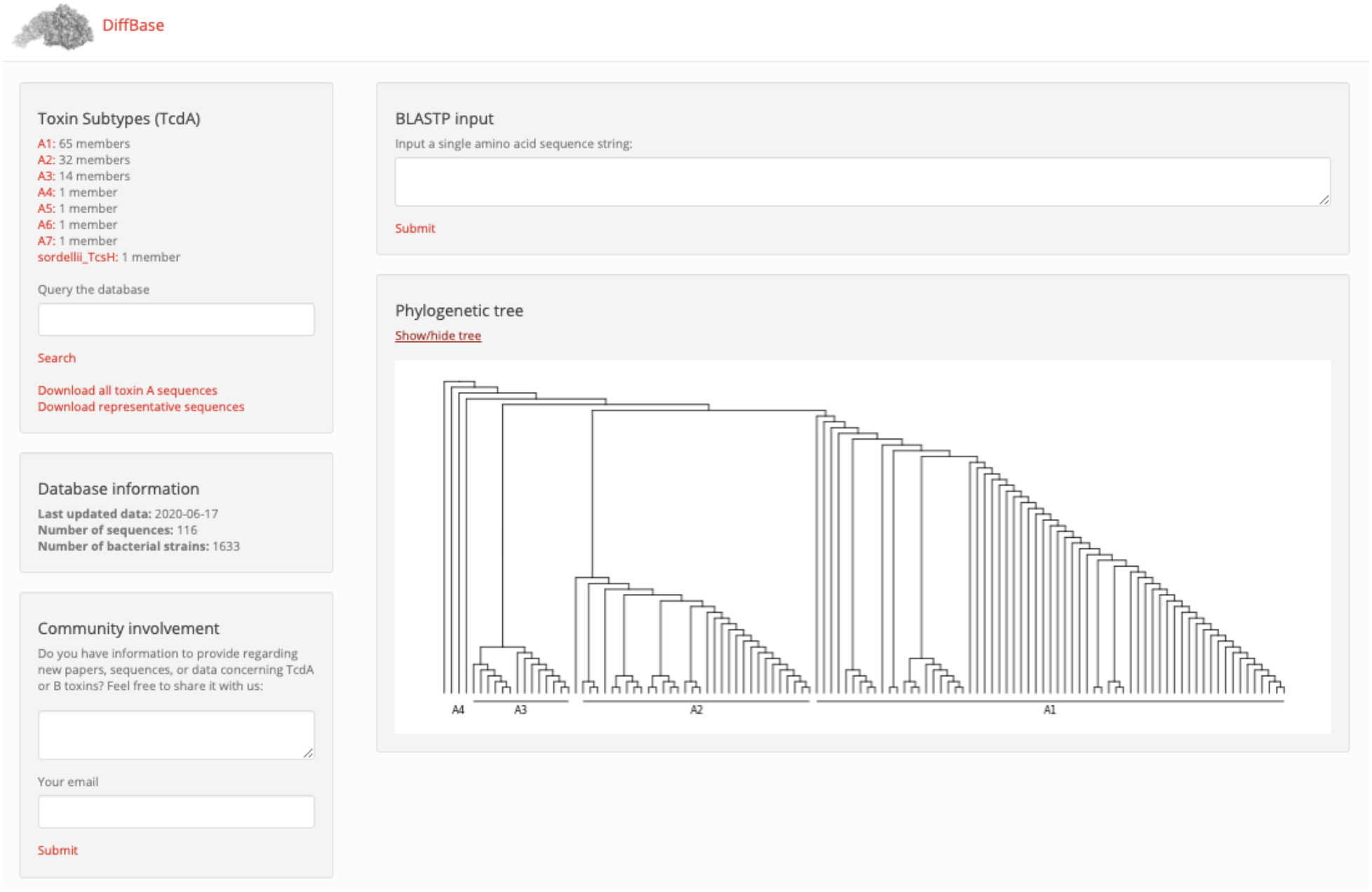
A screenshot of the DiffBase online database, available at diffbase.uwaterloo.ca. DiffBase is currently subdivided into two main sections for TcdA and TcdB sequences. Shown above is the TcdA portion of the database.

**Table S1.** Subtypes of novel TcdA and TcdB sequences identified in the NCBI short read archive. Novel sequences contain at least one substitution not observed in existing sequences derived from NCBI GenBank.

| Subtype | # |
| --- | --- |
| A1 | 25 |
| A2 | 10 |
| A3 | 3 |
| A4 | 1 |
| A5 | 1 |
| A6 | 1 |
| B1 | 52 |
| B2 | 12 |
| B3 | 1 |
| B4 | 7 |
| B5 | 2 |
| B6 | 2 |
| B7 | 1 |
| B8 | 4 |
| B9 | 3 |

**Table S2.**
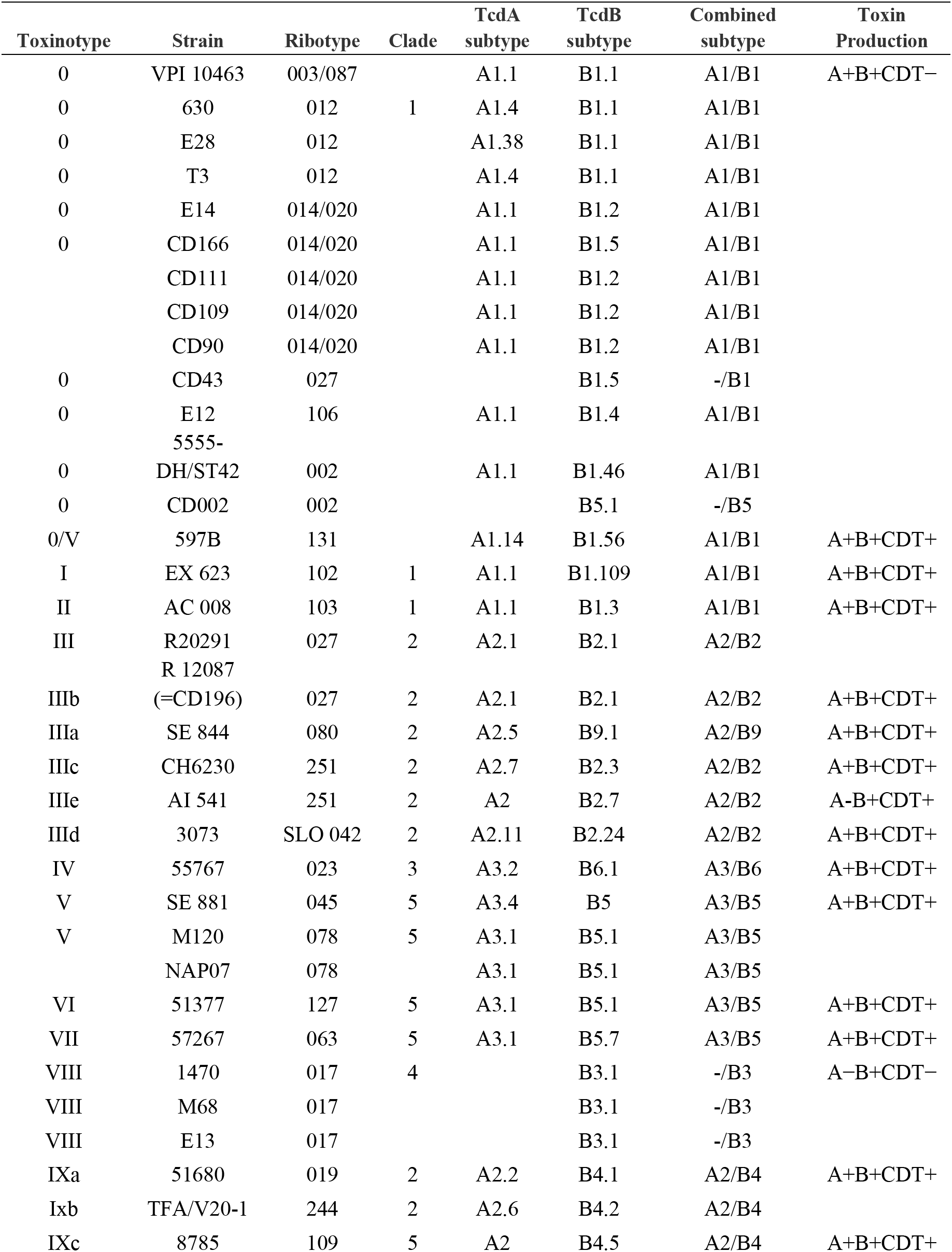
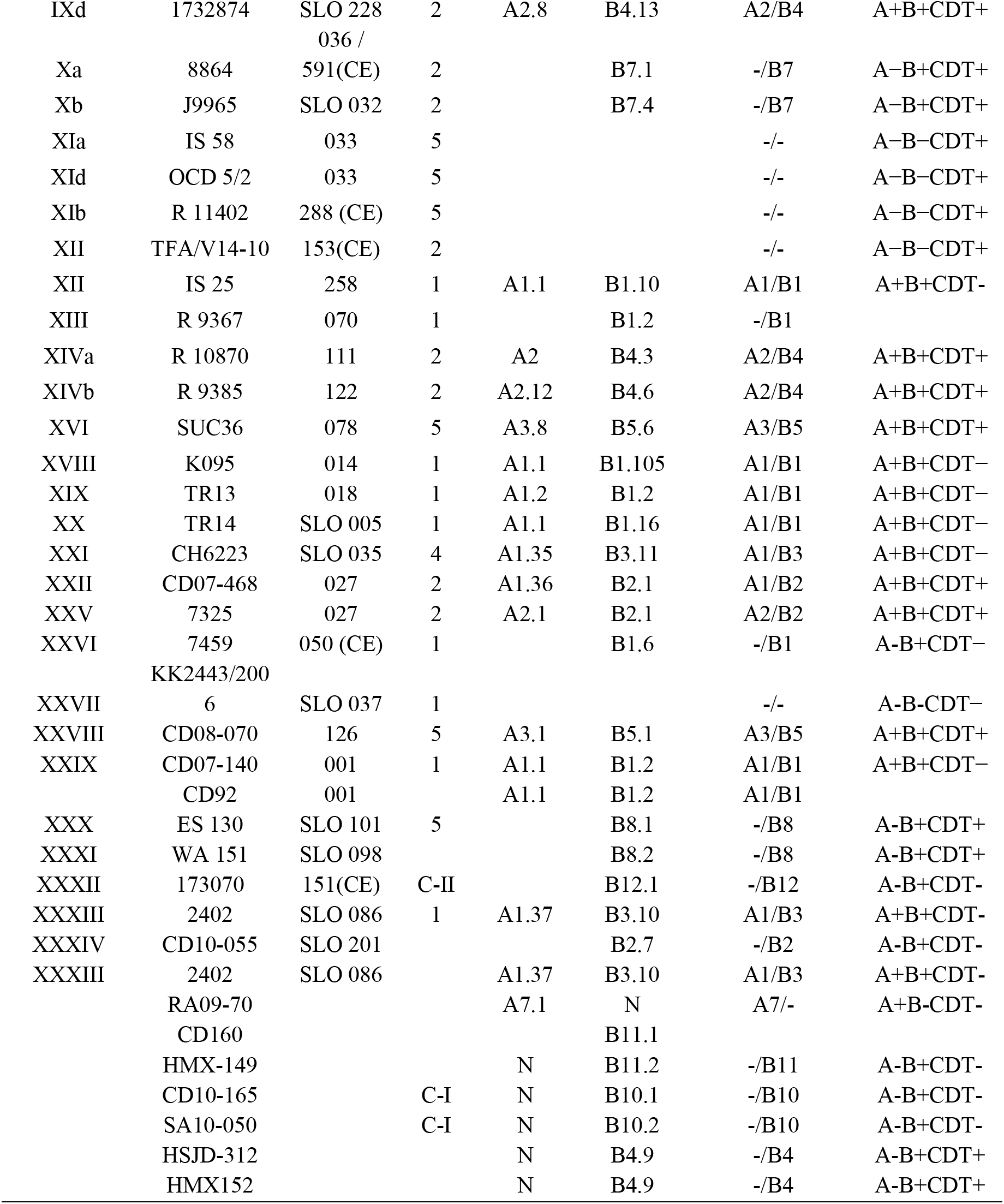
List of clinically relevant and previously studied *C. difficile* strains, associated toxin subtypes, and toxinotypes. Different groups are assigned unique colors. The table is based on information compiled from Rupnik and Janezic (2016), Bletz et al. (2018), and NCBI genome metadata.

